# Lipid oxidation alters membrane mechanics favoring permeabilization

**DOI:** 10.1101/2024.05.23.595481

**Authors:** Sara Lotfipour Nasudivar, Lohans Pedrera, Ana J. García-Sáez

## Abstract

Ferroptosis is a form of regulated necrosis characterized by the iron-dependent accumulation of lipid peroxides in cell membranes. However, how lipid oxidation affects biophysical properties of cellular membranes and how these changes contribute to the opening of membrane pores are major questions in the field. Therefore, understanding the interplay between oxidized phospholipids and alterations in membrane biophysical properties is essential to unravel the underlying mechanisms of ferroptosis. Here, we characterized membrane alterations upon lipid oxidation in *in vitro* model membrane systems using lipid vesicles and supported lipid bilayers (SLBs) in which lipid oxidation was induced by Fenton reactions. We find that vesicle permeabilization kinetically correlates with the appearance of malondialdehyde (MDA), a product of lipid oxidation. Lipid oxidation also alters the lateral organization of SLBs exhibiting liquid-ordered (Lo)/liquid-disordered (Ld) phase coexistence, reducing the lipid mismatch between domains and decreasing the circularity of Lo domains, indicating a decrease in line tension at the phase boundaries. Further characterization of SLBs by force spectroscopy reveals a significant reduction in the average membrane breakthrough force upon oxidation, indicating changes in lipid bilayer organization that make it more susceptible to permeabilization. Our findings suggest that lipid oxidation induces strong changes in membrane lipid interactions and in the mechanical properties of membranes leading to reduced tension in the permeabilized state of the bilayer, which promotes membrane pore formation.

## Introduction

The plasma membrane acts as a permeability barrier that separates the inside from the outside of the cell, thus defining the cell and limiting the transport of molecules. It consists of a closed bilayer of amphiphilic lipids with embedded proteins that together determine its physical properties [1]. Retaining plasma membrane integrity is critical for cellular function and several forms of regulated cell death culminate with plasma membrane disruption. Among them, ferroptosis is a caspase-independent form of regulated necrosis characterized by the accumulation of iron-dependent lipid peroxides within cellular membranes [2]. In contrast to other types of regulated cell death, which possess a specific protein machinery to mediate plasma membrane disruption, no protein executors have been identified that mediate plasma membrane permeabilization or pore formation in ferroptosis. Instead, ferroptosis is induced when lipid oxidation overwhelms cellular antioxidant defenses [2,3]. Ferroptotic cells are characterized by plasma membrane rupture and alteration of membrane phospholipids properties [2]. Among them, the formation of nanopores at the plasma membrane has emerged as a central mechanism for the execution of the final step of this type of cell death [4].

Recent studies have suggested that non-enzymatic lipid peroxidation by Fenton-like reactions are the predominant mechanism driving membrane oxidation during ferroptosis [9,10]. In its simplest form, the Fenton reaction (*H*_2_*O*_2_ + *Fe*^2+^ ➔*Fe*^3+^ +·*OH* + *HO*^-^) describes the reaction between Fe^2+^ and H_2_O_2_ to form a hydroxyl radical (·OH) and a hydroxyl anion [11,12]. In the cellular environment, the hydroxyl radical initiates lipid peroxidation in a chain reaction that depends on the presence of polyunsaturated fatty acids in the membrane phospholipids and on the oxygen concentration, resulting in the formation of peroxidized lipids. Peroxidized lipids tend to hydrolyze into oxidatively truncated oxidized phospholipids with shorter acyl chains which play key role in membrane permeabilization and ferroptosis execution [13]. However, the role of oxidized lipids in the alteration of the biophysical and mechanical properties of membranes that are required for plasma membrane permeabilization in ferroptosis remain poorly understood.

Here, we investigated membrane alterations upon lipid oxidation induced by Fenton-like reactions in two *in vitro* model membrane systems using lipid vesicles and SLB. Lipid model membranes are chemically controlled systems that can be used to study fundamental changes in lipid bilayer organization upon lipid oxidation. Using a liposome leakage assay, we found that vesicle permeabilization kinetically correlates with the appearance of malondialdehyde (MDA), a product of lipid oxidation. We also show that lipid oxidation alters the lateral organization of phase separated membranes, leading to changes in topography and domain shape, by decreasing the line tension between the Lo and Ld phases. Finally, we associate membrane remodelling upon oxidation with a decrease in the average breakthrough force of the membrane. Our results support a general role for lipid oxidation as a promoter of strong changes in the mechanical properties of membranes, which are relevant for membrane pore formation and plasma membrane disruption in ferroptosis.

## Results

### Lipid oxidation increases the permeability of lipid vesicles

Accumulation of oxidized lipids species is recognized as a hallmark of ferroptosis [13–15]. It is also known that lipid peroxidation induced by oxidative stress changes membrane organization and composition resulting in significant alterations such as increased membrane permeability and modifications in packing, fluidity, and viscosity [13,16–19]. To assess whether lipid oxidation directly affects membrane permeability we used a minimal reconstituted system devoid of other cellular components and based on pure lipid, Large Unilamellar Vesicles (LUVs) made of different compositions. We induced lipid oxidation via Fenton-like reactions by adding ascorbic acid and FeSO_4_ to the vesicle mixtures (Figure 1A). Vesicle permeability changes were assessed by monitoring the release of carboxyfluorescein (CF), a self-quenching dye encapsulated within the LUVs (Figure 1A). Using this liposome leakage assay, we found that incubation with iron and ascorbic acid lead to permeabilization LUVs made of egg-PC whereas neither ascorbic acid nor FeSO_4_ individually were able to affect vesicle stability (Figure. 1B). We also observed a dose-dependent relationship between ascorbic acid concentration and CF release from LUVs made of egg-PC, with maximal permeabilization capacity at ascorbic acid concentrations above 2.4 mM (Figure. 1C). To characterize the temporal relationship between lipid oxidation and LUVs permeabilization, we tracked in parallel the carboxifluorescein fluorescence and lipid oxidation measured by the thiobarbituric acid reactive substance (TBARS) assay (Figure 1D). The appearance of malondialdehyde (MDA), a secondary product of lipid oxidation, kinetically correlated with vesicle permeabilization (Figure 1C). Egg-PC vesicles exhibited a permeabilization extent of about 80% of CF release, in contrast to vesicles made of DOPC or DOPC:SM:Chol (2:2:1), which were little permeabilized by incubation with the same concentrations of iron and ascorbic acid. These results suggest that the polyunsaturated fatty acids present in egg-PC make the lipid bilayers more susceptible to oxidation compared to the monounsaturated fatty acids present in DOPC or in SM (Figure 1B). Further increases in the concentrations of ascorbic acid and FeSO_4_ were required to permeabilise DOPC vesicles up to 30 % of CF release (Figure 1F). Incorporation of saturated SM as well as Chol into DOPC vesicles decreased the permeabilizing activity induced by lipid oxidation, which can be related to the decrease in the proportion of lipids with unsaturated fatty acids (Figure 1F). To characterize the effect of lipid phase coexistence on the sensitivity of LUVs to lipid oxidation, we used vesicles made by the ternary mixture DOPC:SM:Chol (2:2:1) which presents Lo/Ld phase coexistence (Figure 1E). Collectively, our experiments show that lipid oxidation induced by Fenton-like reactions causes permeabilization of lipid vesicles of different compositions, confirming that the presence of oxidized lipid species is sufficient to induce membrane permeabilization.

**Figure 1.**
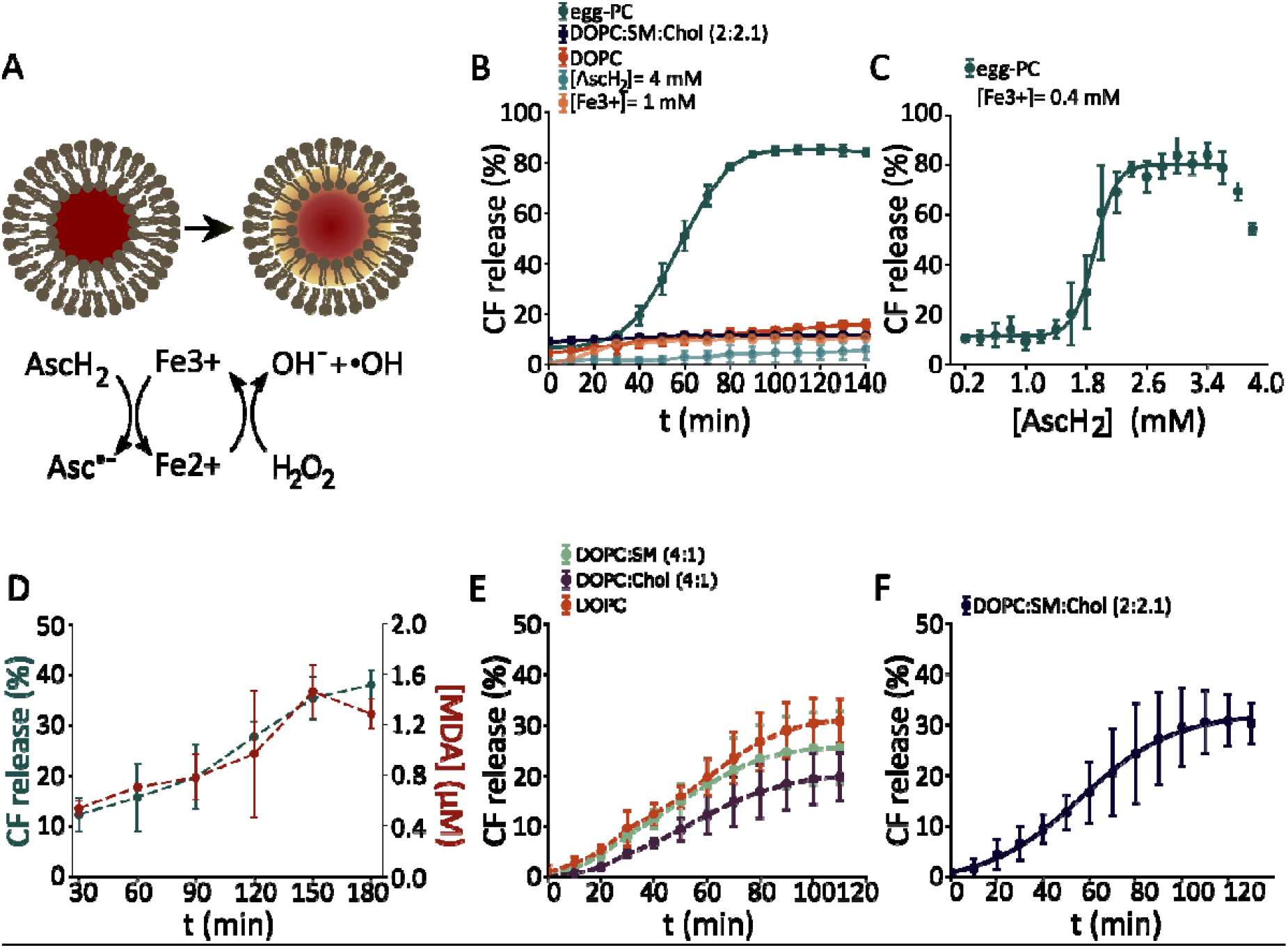
Lipid oxidation induces permeabilization of lipid vesicles. (A) Schematic representation of the experimental setup. Iron sulphate and ascorbic acid initiate Fenton-like chain reactions of lipid oxidation. Permeabilization of lipid vesicles increases the leakage of carboxyfluorescein. (B) Kinetic of carboxifluorescein leakage from liposomes made of eggPC, DOPC or DOPC:SM:Chol (2:2:1) upon lipid oxidation induced by 0.8 mM iron sulfate and 2 mM ascorbaic acid at 37°C. The concentrations of iron sulphate and ascorbic acid used for negative control experiments are indicated on the figure. (C) Dosis-response curve of permeabilization of LUV made of eggPC as a function of ascorbic acid concentration in the presence of 400 μM iron sulphate. (D) Kinetic curves of vesicle permeabilization and malondialdehyde concentration in egg-PC vesicles treated with 400 μM iron sulphate and 1.6 mM of ascorbic acid. (E) Kinetics of permeability increase in vesicles made of DOPC, DOPC:SM (4:1) and DOPC:Chol (4:1) upon lipid oxidation induced by 800 μM iron sulphate and 8 mM ascorbic acid. (F) Kinetic of permeabilization in vesicles made of DOPC:SM:Chol (2:2:1) treated with 8 mM ascorbic acid and 500 μM iron sulphate. All data points represent mean values and bars indicate the standard deviations from a set of at least three independent experiments. When no error bar is observed the corresponding standard deviation is smaller than the size of the symbol.

### Lipid oxidation affects the lateral organization of the lipid membrane

Lipid oxidation has been observed to shift the miscibility of components in ternary lipid systems, resulting in the appearance of phase coexistence in a previously homogenous membrane [19]. To investigate whether lipid oxidation affects the lateral organization of membranes, we visualized supported lipid bilayers (SLBs) made of DOPC:SM:Chol (2:2:1) exhibiting Lo/Ld phase coexistence by confocal imaging and by atomic force microscopy (AFM) (Figure 2). To visualize lipid peroxidation in SLBs, we used the lipid peroxidation sensor C11 BODIPY 581/591. In mixtures with Lo/Ld phase coexistense, Lo domains exhibit round shapes that tower over the Ld continuos phase due to a height mismatch [20– 24]. Under these conditions, the hydrophobic tails of the lipids minimize exposure to water by tilting, which causes a line tension at the domain edge.

**Figure 2.**
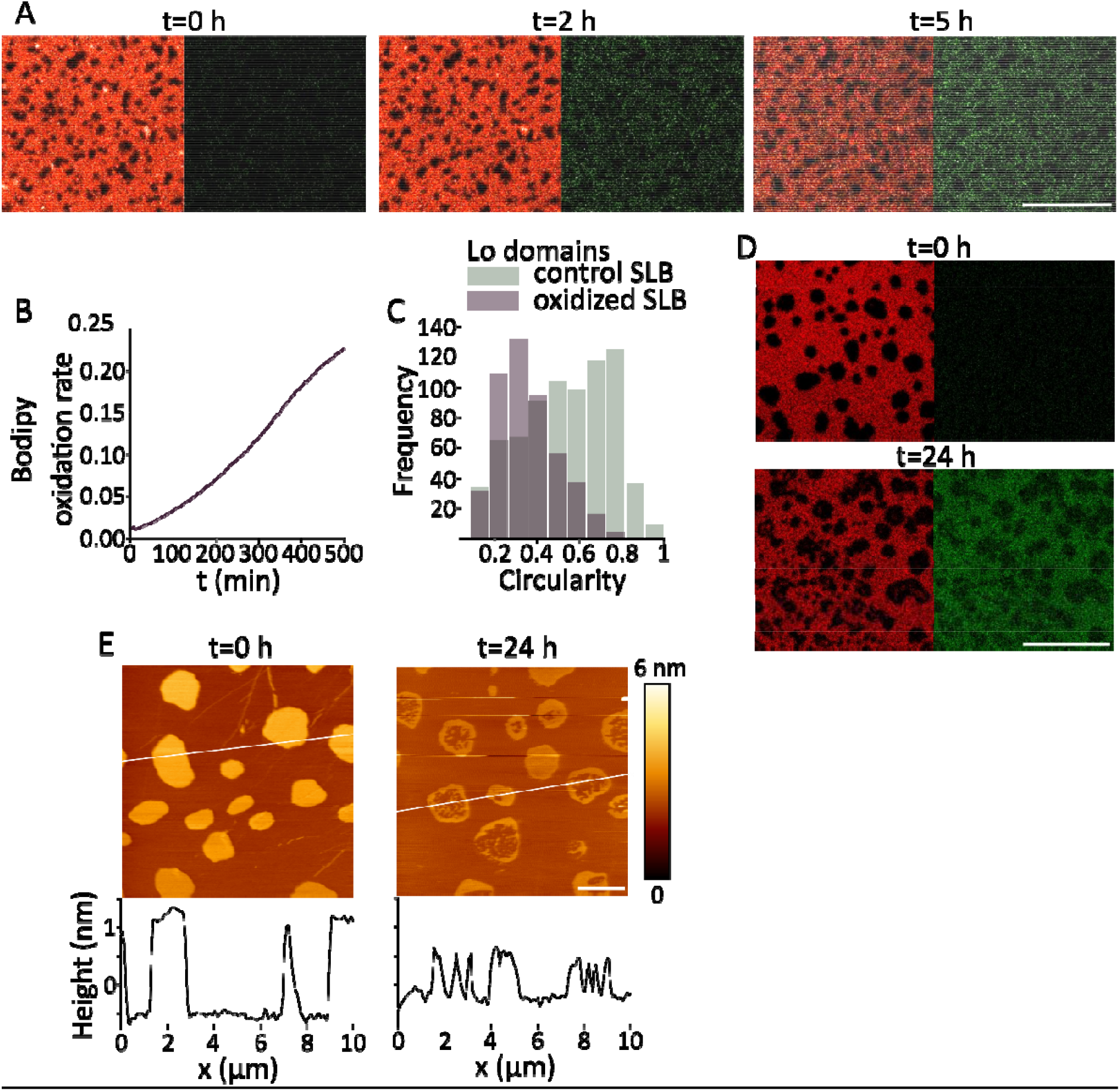
Changes in membrane lateral organization during oxidation. (A) Confocal images of SLBs with composition DOPC:SM:Chol (2:2:1) and labelled with the fluorescent dye BODIPY-C11 (0.1 mol%), before and after 2 and 5 h incubation with 8 mM ascorbic acid and 500 μM iron sulfate at 37 °C. (B) Time course of the increase in the normalized oxidation ratio. (C) Quantification of the circularity of Lo domains in untreated and oxidized SLBs after 24 h of treatment. (D) Corresponding confocal images of SLBs stained with BODIPY-C11 (0.1 mol%) before oxidation and at 24 h, showing changes in domain shape. Scale bars, 10 μm. (E) AFM images of SLBs made of DOPC:SM:Chol (2:2:1) after 5 h oxidation induced with 8 mM ascorbic acid and 500 μM iron sulphate at 37 °C. Corresponding line profiles as indicated by white line in AFM image above, t = 0 h corresponds to a non-oxidized SLB with Lo and Ld domains. Scale bar, 2 μm.

We found that incubation with iron and ascorbic acid promoted lipid peroxidation at the SLBs detected an increase in the green fluorescent signal of BODIPY starting at 5 hours (BODIPYox) (Figure 2A and B). Lipid oxidation altered membrane topography decreasing the circularity of Lo domains (Figure 2C,D). The precise characterization of the SLBs using AFM additionally revealed a decrease in the lipid mismatch height between domains upon oxidation (Figure 2E). These two changes indicate a decrease in line tension at the phase boundaries. Interestingly, we also detected the appearance of Ld within Lo domains, suggesting a reduction of the average lipid order in the SLBs (Figure 2E).

### Lipid oxidation affects the fluidity of supported lipid bilayer

To further study the effect of lipid oxidation on membrane fluidity, we performed experiments of fluorescence recovery after photobleaching (FRAP) (Figure 3). In non-oxidized bilayers, the measurements were made only in the Ld phase whereas in the oxidized bilayer this was not possible due to the small size of the Lo domains (Figure 3B). Interestingly, lipid oxidation caused a decrease in the immobile membrane fraction and an increase in the fluorescence recovery kinetics in comparison with non-oxidized bilayers (Figure 3). These results suggest that lipid oxidation causes reorganization of the lipid bilayer into a less compartmentalized environment with increased movement of the lipid components, in line with the remodelling of SLBs observed by confocal microscopy and AFM (Figure 2).

**Figure 3.**
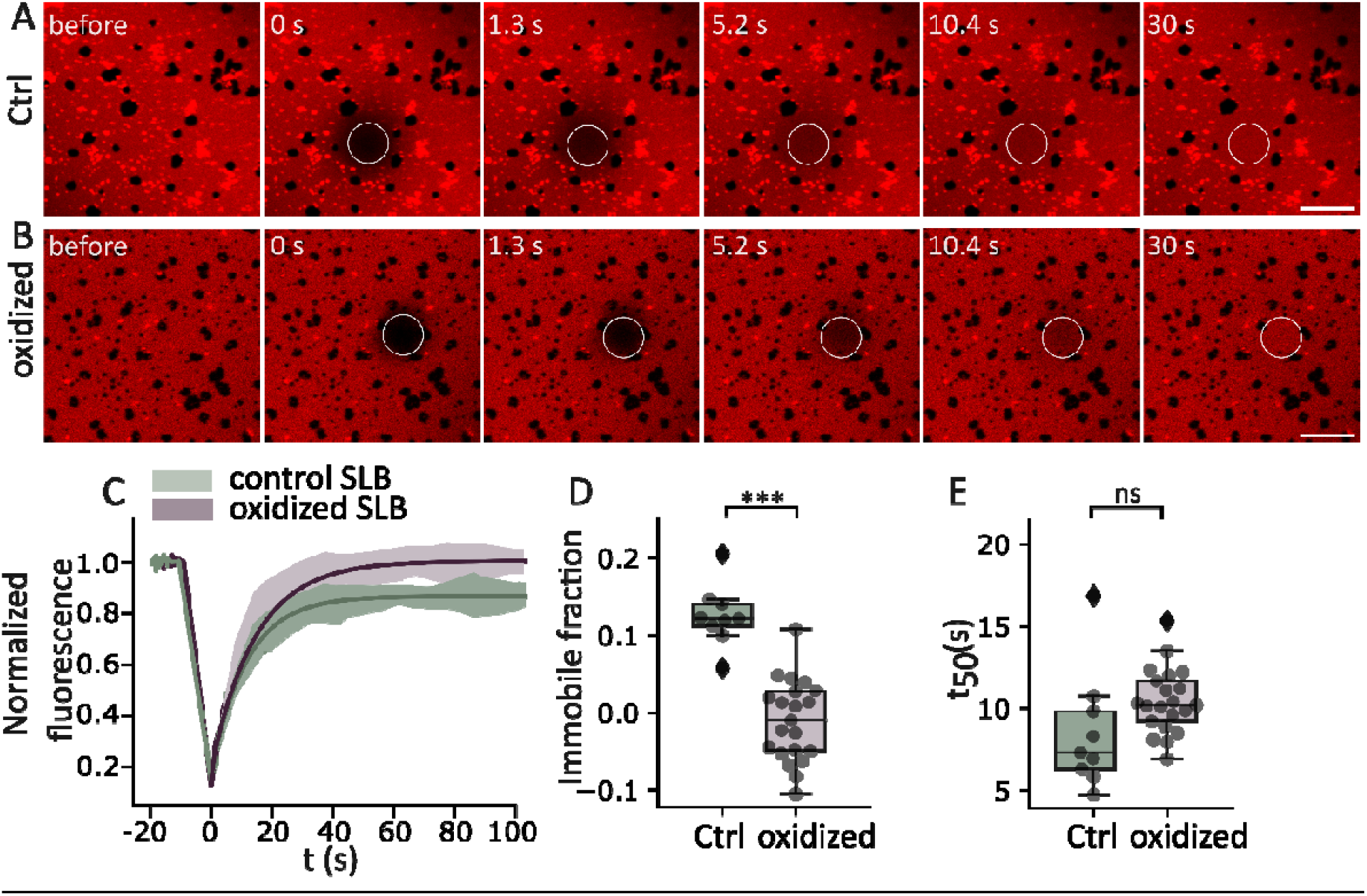
Lipid oxidation increases the lateral movement of lipids in SLBs. Time-lapse confocal images of a representative FRAP experiment in SLBs made of DOPC:SM:Chol (2:2:1) and labelled with rhodamine-PE (0.5 mol%) in (A) non-oxidized bilayer, (B) oxidized bilayer after 24 h treatment with 8 mM ascorbic acid and 500 µM iron sulphate. Scale bars, 10 μm. (C) FRAP recovery curves in SLBs before and after lipid oxidation. The graph shows the average (n = 9 individual curves in the non-oxidized bilayers and n = 21 individual curves in oxidized bilayers) temporal increase of the normalized fluorescence. Immobile fraction (D) and t_50%_ (E) are calculated by fitting the corresponding recovery curves to an exponential function. Each dot represents a single measurement. In box plots (D and E), the line inside the box indicates the median, the box indicates the interquartile range (IQR) of the values, and the error bars indicate the 1.5 IQR. Statistical comparison between groups was performed by Student’s t-test, *** indicates independent groups with significant differences p < 0.001.

### Lipid oxidation decreases the breakthrough force of the membrane

In addition to the high-resolution imaging of SLBs topography, AFM also allows the determination of the physical properties of the membrane in membrane piercing experiments using force spectroscopy (FS). By calibrating the cantilever, its deflection can be used to determine force values. Typical approaching FS curves of oxidized and non-oxidized SLBs made of DOPC:SM:Chol (2:2:1) are shown in Figure 4B. As the AFM tip contacts and pushes against the membrane surface, the force gradually increases until a critical threshold is reached, at which point the membrane is punctured, resulting in a sudden drop in force (breakthrough force).

**Figure 4.**
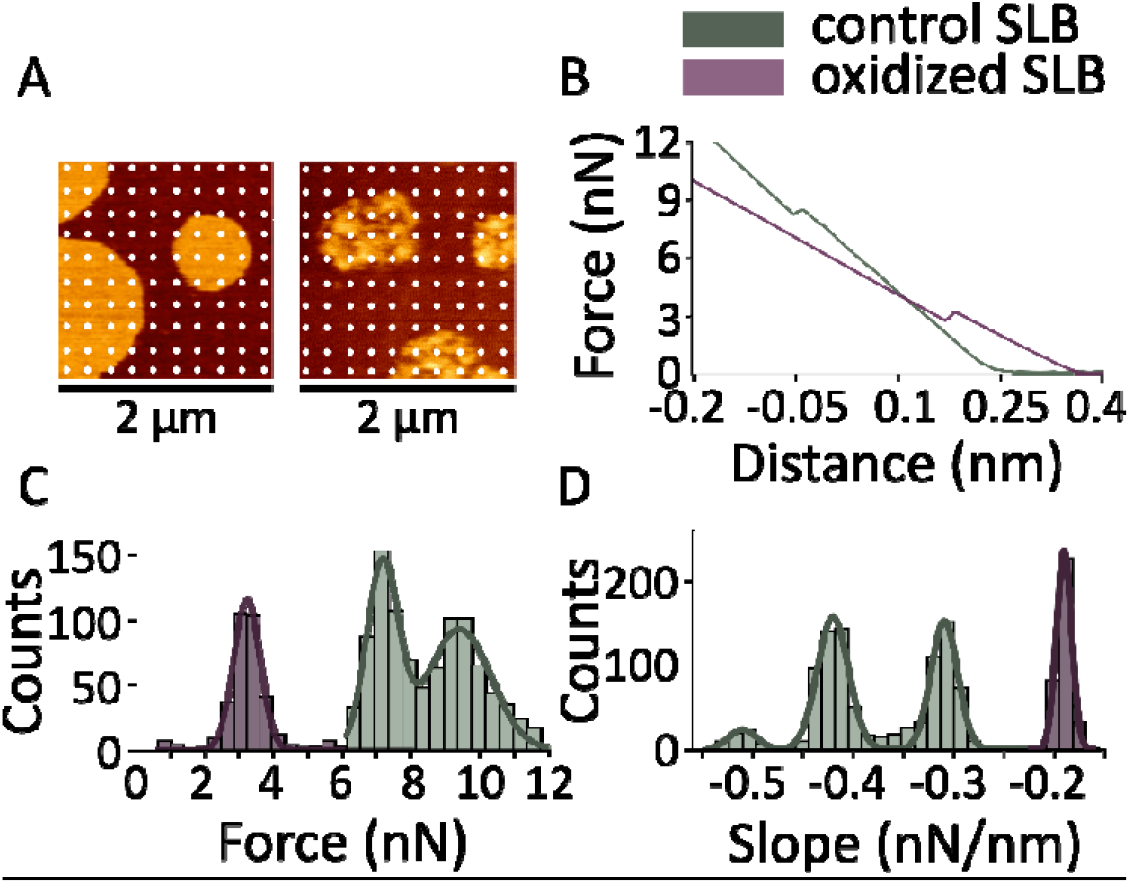
Oxidation decreases the force needed to pierce the membrane. (A) Example grid of force measurements on non-oxidized (left) and oxidized SLBs (righ) made of DOPC:SM:Chol (2:2:1) after 24 h treatment with 8 mM ascorbic acid and 500 µM iron sulphate. Each dot represents a position where a force spectroscopy measurement was performed. Images 2 µm x 2 µm. (B) Representative force spectroscopy curves for non-oxidized and oxidized bilayers. Histograms for the distribution of piercing forces (C) and force/distance slopes (D) for non-oxidized (n=954) and oxidized bilayer (n=343). Lines are the best fit of the histograms data to bi- and trimodial Gaussian for determination of mean and standard deviation (R2 > 0.97).

We found that untreated bilayers exhibit two populations of breakthrough forces around 9.4 nN ± 0.9 nN and 7.2 nN ± 0.5 nN corresponding to the forces required to puncture the liquid-ordered (Lo) and liquid-disordered (Ld) phases, respectively (Figure. 4C). After 12 hours of incubation with iron and ascorbic acid, we observed that membrane remodelling is accompanied by a significant reduction of the average breakthrough force to 3.2 nN ± 0.4 nN and the loss of the bimodal behaviour, indicating the mixing of lipids from the two liquid phases (Figure 4A and C). Non-oxidized SLBs showed two populations of the slope values around -0.31 ± 0.01 nN/nm and -0.42 ± 0.02 nN/nm whereas the oxidized bilayer showed a single population around -0.19 ± 0.01 nN/nm (Figure 4D). Overall, our results show that lipid oxidation induces strong changes in the mechanical properties of the membrane, leading to a decrease in membrane tension in correlation with the permeabilized state of the bilayer.

## Discussion

Despite the rapid progress in understanding ferroptosis in recent years, how lipid oxidation affects membrane biophysical properties and how these changes contribute to membrane pore opening remain major questions in the field. Here, we investigated membrane alterations upon lipid oxidation induced by Fenton-like reactions in *in vitro* model membrane systems using lipid vesicles and SLBs. We found that lipid oxidation increases liposome permeability, which is associated with increased membrane fluidity and reorganization of lipid domains, leading to a reduction in the liquid order phase area and in the height mismatch between the two phases. Interestingly, we also detected temporal changes in membrane mechanics towards reduced tension in the permeabilized state.

Alteration of plasma membrane permeability is a hallmark of ferroptosis that is common to other types of regulated necrosis such as necroptosis and pyroptosis. In contrast to necroptosis and pyroptosis, in which membrane permeabilization is mediated by protein executors, oxidized lipids are thought to be directly responsible for plasma membrane breakdown in ferroptosis. During oxidation, lipid hydroperoxides are initially formed and later truncated to their derivative oxidatively truncated phospholipid species [25,26]. Structural modifications of oxidized species have been shown to disrupt the regular packing of lipids within the bilayer, associated with changes in membrane integrity and permeability [26–29]. Truncated oxidized lipids, such as PazePC, can play an important role in membrane permeabilization, even at low concentrations [29]. Their conical shape and ability to induce positive membrane curvature contribute to the formation of pores and defects in the membrane [30,31]. Recently, the key role of oxidatively truncated phospholipid species in membrane permeabilization and ferroptosis execution has been presented [13].

We also found that lipid oxidation induces the lateral reorganization of lipid domains in SLBs, reducing the lipid mismatch between the Lo and Ld lipid phases. The increased polarity of lipid hydroperoxides, and especially oxidatively truncated lipids, affects their interactions with neighbouring lipid molecules, potentially affecting the overall organization of the membrane. Our FRAP experiments indicate an altered diffusion behaviour that supports this hypothesis. Changes in shape and length of oxidized species induce a more heterogeneous bilayer and could lead to disruption of the regular packing. Furthermore, molecular dynamics simulations suggested that cholesterol and PazePC can colocalize and form nano domains [28]. Under our experimental conditions, the colocalization of oxidized lipids like PazePC and cholesterol could possibly lead to a reduction in the size of the Lo domains, which would decrease the immobile fraction.

In any case, our results indicate that the membrane becomes more susceptible to mechanical perturbations, leading to a decreased resistance to puncture, making fluid membranes more susceptible to deformation and disruption under external forces. The relationships between Young’s modulus, force and penetration depth [33] allow us to compare the slope values and we further conclude that oxidized membranes present decreased rigidity. The loss of the bimodal behaviour in both the breakthrough forces and the slopes of the force-distance curves strongly suggests that oxidized lipid species induce alterations in mechanical properties that affect broadly throughout the bilayer.

In summary, we found here that lipid oxidation induced by Fenton-like reactions alters the lateral organization of lipids in bilayers and the mechanical properties of the membrane, resulting in the loss of its permeability barrier function. We propose that oxidized lipids, by altering lipid packing, reduce the high energetic cost of pore formation in the lipid bilayer, which may be of relevance in the context of ferroptosis. Pore opening would then allow water entry and further affects the mechanical properties of the bilayer prior to membrane rupture.

## Materials and Methods

### Reagents

All reagents and lipid standards were purchased from Sigma Aldrich (St. Louis, USA) or Avanti Polar Lipids (Alabaster, USA). Organic solvents were purchased from Supelco/Merck KGaA (Darmstadt, Germany). The oxidation of liposomes was induced by the addition of various concentrations of FeSO_4_ and ascorbic acid and incubated at 37°C at the indicated times.

### Permeabiliziation Assay of large unilamellar vesicles

L-α-phosphatidylcholine from egg yolk (egg-PC) was dissolved in carboxyfluorescein (CF) (80 mM, pH 7.0) to a concentration of 5 mg/mL. After six cycles of freezing and thawing, the lipid mixture was extruded by 31 passes through a 100 nm pore size membrane. A Sephadex G50 column was used to separate liposomes with encapsulated CF from the non-encapsulated CF using as outside buffer 300 mM NaCl, 20 mM Hepes pH7.0. A ratio of fluorescence between permeabilized and non-permeabilized vesicles greater than 3 was used. Treatment of the vesicles with iron and ascorbate at different concentrations initiates a Fenton reaction and leads to lipid peroxidation of the phospholipids. The fluorescence signal corresponding to CF was measured every 10 min with a plate reader (Perkins) using an excitation wavelength of 485 nm and an emission wavelength of 525 nm. We used Triton X-100 to induce the total permeabilization of the vesicles in the assay. In parallel, we measured the fluorescence of untreated vesicles as a reference for non-permeabilized vesicles. The percentage of permeabilized vesicles was calculated as follows: 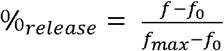

### Measurement of Malondialdehyde with the Thioarbituric Acid Assay

The thioarbituric acid (TBA) assay measures the reaction product of malondialdehyde (MDA) and TBA and can be used as a reference for the level of oxidation. MDA is a secondary oxidation product formed during lipid peroxidation. Two molecules of TBA form a chromogen that allows the spectrophotometric determination at 532 nm [32].

### Supported Lipid Bilayer

SLBs were prepared for AFM measurements and confocal microscopy. Lipid compositions were prepared and stored as a dry film. The lipids di-oleoyl-phosphatidylcholine (DOPC), sphingomyelin (SM) and cholesterol (Chol) were dissolved in chloroform. The lipid film was rehydrated in PBS buffer (2.7 mM KCl, 1.5 mM KH_2_PO_4_, 8 mM Na_2_HPO_4_, 137 mM NaCl, pH 7.2) to a final concentration of 10 mg/mL and stored at -20°C. The lipid mixture was resuspended in 140 *µ*l of buffer (10 mM HEPES, 150 mM NaCl, 3 mM NaN_3_, pH 7.2) and sonicated for 15 min to form SUVs. The lipid suspension was deposited to a freshly prepared mica disk on a glass slide and CaCl_2_ was added to a total concentration of 3 mM in a final volume of 500 *µ*l. After an initial incubation of 2 minutes at 37°C followed by 10 minutes at 65°C and slow cooling, floating vesicles and CaCl_2_ were washed out by adding and removing 150 *µ*l of buffer solution 20 times. SLBs were additionally stained with Bodipy BODIPY-C11 (0.1 mol%) for the tracking of oxidation or labelled with rhodamine-PE (0.5 mol%) for FRAP experiments.

### Confocal microscopy

Confocal fluorescence microscopy was performed using an Olympus infinity line scanning microscope (Abberior Instruments) equipped with a UPlanX APO 60x Oil/1.42 NA objective. Lo circularity 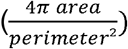 values were analyzed using Image J (http://imagej.nih.gov/ij/) analyze particles function. Photobleaching experiments were performed using 60x Oil 8x8 *µ*m area was bleached at nominal 100% laser transmission and a serious of 5 images, following immediate recovery was captured for a serious of 80 images (interval 1.2 s).

### Atomic Force Microscopy - Imaging

Lipid bilayers were imaged using a JPK nanowizard (JPK Instruments, Berlin, Germany) in contact or tapping mode and V-shaped non-conducting silicon nitride cantilevers (Bruker, DNP-10) with a typical spring constant of 0.06 Newton/m. The scanning rate was set between 0.6 and 2 Hz. Images were processed by applying a degree 1 or 2 polynomial fit to the surface.

### Atomic Force Microscopy - Force Spectroscopy

For this measurement, an area was selected on a bilayer topographical image on which a set of 10x10 or 14x14 force measurements were operated. A limit is set at which the cantilever is lowered at a constant speed. The dependence of the force as the function of height can be plotted using Python. Before generating the force histograms, a peak detection script was run in Python given out a file with detected values.

## Acknowledgements

This work was supported by the Deutsche Forschungsgemainschaft, grant n. GA1641/7-1 from the program SPP2306 “Ferroptosis: from basic research to clinical applications”, and by the University of Cologne.

